# “Hemicellulose Degradation and Utilization by a Synthetic *Saccharomyces cerevisiae* Consortium”

**DOI:** 10.1101/244244

**Authors:** Ian Dominic F. Tabañag, Shen-Long Tsai

## Abstract

Since *Saccharomyces cerevisiae* does not inherently possess the capability to utilize pentose sugars released from hemicellulose degradation, the degradation and utilization of hemicellulose poses a conundrum to bioethanol production by consolidated bioprocessing (CBP) using *S. cerevisiae*. In this study, *S. cerevisiae* was exploited for its ability to degrade xylan, one of the major polysaccharide chains present in hemicellulose. Different hemicellulases from *Trichoderma reesei*, namely: endoxylanase (Xyn2), *β*-xylosidase (Bxl1), acetylxylan esterase (Axe1), *α*-D-glucuronidase (Glr1) and *α*-L-arabinofuranosidase (Abf1), were heterologously secreted by *S. cerevisiae*. A mixture experimental design was adapted to statistically describe the synergistic interactions between the hemicellulases and to determine the optimum formulations for the hydrolysis of xylan substrates. The hydrolytic activities of the hemicellulase mixtures were then improved by displaying the hemicellulases on the yeast surface to serve as whole-cell biocatalysts. The engineered yeast strains displaying hemicellulases were further engineered with xylose-utilization genes to enable abilities of utilizing xylose as a sole carbon source. The resulting consortia were then able to grow and produce ethanol from different xylan substrates.

## 1 Introduction

The most abundant biomass found in nature which is preferably exploited in consolidated bioprocessing (CBP) is lignocellulose. Because of its abundance, this raw material can be obtained in fact at a very low cost, making it the best substrate with the highest potential in CBP. Moreover, the cellulosic ethanol obtained from lignocellulosic substrates has been reported sustainable as its production process has reduced greenhouse gas emissions (Langeveld, Sanders et al. 2012, Salles-Filho, Cortez et al. 2016). Lignocellulose comprises 50–90% of all plant matter and it is composed of three major components namely: cellulose (~35–50%), hemicellulose (20–35%; which is composed mainly of xylan), and lignin (~15–30%). These major components of lignocellulose are in their polysaccharide forms and hydrolysis of these polysaccharides yield fermentable sugars except for lignin (which produces phenolic compounds that actively inhibit fermentation (Lynd, Weimer et al. 2002, Kricka, Fitzpatrick et al. 2014).

Hemicellulose is mainly composed of pentose sugars such as: xylose, arabinose and mannose. One of the most challenging conundrums to use *S. cerevisiae* in CBP is the inability of *S. cerevisiae* to completely break down hemicellulose due to its structural complexity and to utilize the pentose sugars released upon hemicellulose degradation. Owing to the structural complexity of hemicellulose, a mixture of the various hemicellulases which are classified into: main-chain, and side-chain (or known as accessory) enzymes are needed for its total degradation as schematically illustrated in Figure 1 for the efficient degradation and xylose release from the xylan backbone. The interaction of these main-chain and accessory enzymes is described by performing synergy studies (Nidetzky, Steiner et al. 1994, Kumar and Wyman 2009). A given argument between these two types of hemicellulases is that the main-chain cleaving hemicellulase will have an enhanced activity if the substituents (minor components such as arabinose, mannose, and some functional groups such as acetate) are prior removed by the accessory enzymes provided the steric hindrance to the main-chain enzyme of the substituent (Collins, Gerday et al. 2005, Meyer, Rosgaard et al. 2009, Van Dyk and Pletschke 2012, Bhattacharya, Bhattacharya et al. 2015, Moreira and Filho 2016). Van Dyk and Pletschke (2012) provided a good review on the synergistic studies of different lignocellulose-degrading enzymes.

**Figure 1.**
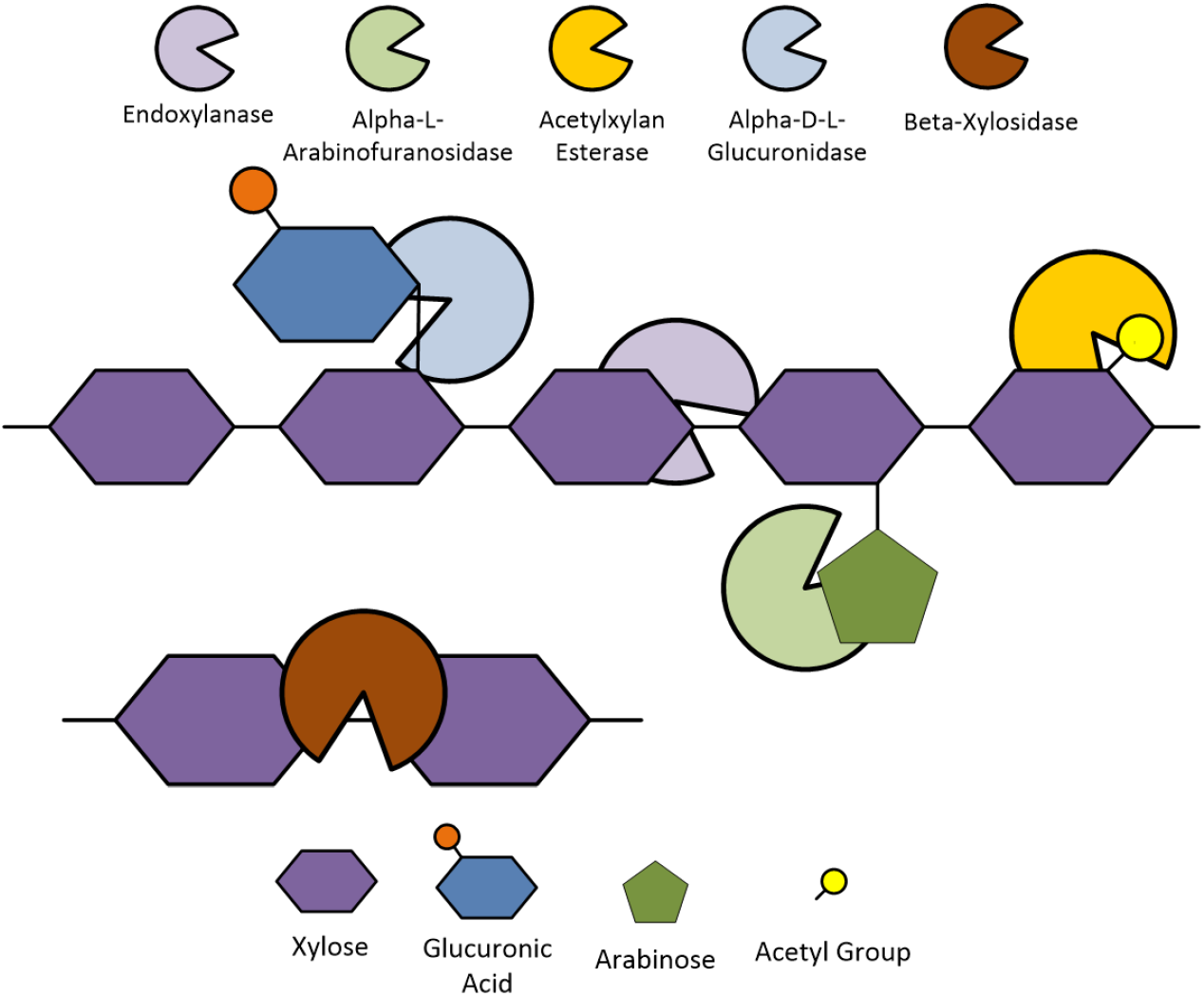
The basic structural components found in hemicellulose and the hemicellulases responsible for their degradation.

Specifically for hemicellulose, a number of synergistic studies have been performed using hemicellulase mixtures purified from different microorganisms. It was generally reported that accessory enzymes such as xylan esterases, arabinofuranosidases, glucuronidases, and mannases synergistically enhances the main-chain cleaving enzymes (i.e., xylanases, xylosidases) in the release of xylose from pure hemicellulose substrates, such as wheat arabinoxylan, oat-spelt, beechwood, and birchwood xylans (Gasparic, Martin et al. 1995, Christakopoulos, Katapodis et al. 2000, Sørensen, Meyer et al. 2003, Vardakou, Katapodis et al. 2004, Kam, Jun et al. 2005, Sørensen, Pedersen et al. 2007, Sørensen, Pedersen et al. 2007, Raweesri, Riangrungrojana et al. 2008, Shi, Li et al. 2010, McClendon, Shin et al. 2011, Dyk and Pletschke 2013, Goldbeck, Damásio et al. 2014). Though it should be noted that not all hemicellulase combinations would lead to a positive synergism between them. Some exhibit negative synergism (also referred as anti-synergism) in cases when one of the enzymes in the mixture inhibits the action of its other constituents (Kovacs 2009). Furthermore, some cases show that some accessory enzymes exhibit no synergy with respect to the main-chain cleaving enzymes; in other words, the addition of the accessory enzyme did not even improve the xylose, arabinose, mannose, or ferulic acid release either due to the nature of the enzyme (e.g, substrate specificity, reaction conditions, etc.) or the complexity of the substrate (de Vries, Kester et al. 2000, Adelsberger, Hertel et al. 2004, Vardakou, Katapodis et al. 2004, Selig, Adney et al. 2009).

However, synergistic studies for different hemicellulase mixtures when expressed in *S. cerevisiae* for CBP are quite limited (La Grange, Pretorius et al. 2001, Srikrishnan, Chen et al. 2013). Thus, this study aims to investigate the following: i.) synergistic action of different types of hemicellulases to xylan substrates, from the hemicellulose-degrading fungus *Trichoderma reesei*, when expressed and secreted in the CBP-microorganism *Saccharomyces cerevisiae* using a statistical mixture experimental design approach, ii.) enhancement of hemicellulase mixture activities via a whole-cell biocatalyst approach iii.) the degradation and utilization of the said xylan subsrates by a mixed culture fermentation approach of the hemicellulase-expressing, xylose-utilizing yeast strains The adaptation of these approaches would lead to the creation whole-cell biocatalysts that have the ability of producing ethanol directly from xylan substrates.

## 2 2. Materials and Methods

### 2.1 Strains and Media

The filamentous fungus *Trichoderma reesei* QM6a, acquired from the Bacterial Culture and Resource Center in Taiwan and cultivated in Potato dextrose broth for maintenance and Mandel’s medium (Mary and James 1969) for hemicellulase expression, was utilized as an mRNA source for the cloning of hemicellulases.

The *Escherichia coli strain* JM109 [*endA1 glnV44 thi-1 relA1 gyrA96 recA1 mcrB^+^ Δ(lac-proAB)* 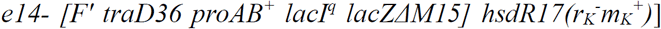, was utilized as a host strain for recombinant DNA manipulation and cultivated in Luria-Bertani medium (1% tryptone, 0.5% yeast extract, and 1% NaCl) supplemented with appropriate antibiotics.

The *Saccharomyces cerevisiae* strain EBY100 *[MATa AGA1::GAL1-AGA1::URA3 ura3–52 trp1 leu2-delta200 his3-delta200 pep4::HIS3 prb11.6R can1 GAL*], was exploited to express and secrete heterologous hemicellulases from *T. reesei;* and cultivated in either YPD Medium (2% dextrose, 1% yeast extract and, 2% peptone) for maintenance or Synthetic Minimal Medium with Casamino Acids (SMD+CAA; 2% glucose, 0.67% yeast nitrogen base and, 0.5% Casamino Acids; SMD (ade+, trp^+^ or leu^+^) for selection.

For ethanol fermentation from hemicellulase substrates, yeast cells were pre-cultivated with either SMD+CAA or SMD media. After pre-cultivation, the yeast cells were then inoculated into SM+Xylan (1 or 2% w/v) or Arabinoxylan (1or 2% w/v) buffered with 50mM Na-citrate (~pH 6) containing 12mM *CaCl_2_* and 2mM EDTA.

### 2.2 Molecular Biology Techniques

Total RNA was isolated from *T. reesei* cultures by Trizol extraction using the Direct-Zol™ RNA Mini-prep Kit (Zymo Research, CA, USA) according to the manufacturer’s specification. cDNA was generated from the total RNA samples by reverse transcription PCR using the PrimeScript™ High Fidelity RT-PCR Kit (TaKaRa Bio Inc., CA, USA) according to the manufacturer’s specification. The five hemicellulases namely: i.) Arabinofuranosidase GH54 (Abf1), ii.) Acetylxylan esterase CE5 (Axe1), iii.) *β*-xylosidase GH3 (Bxl1), iv.) *α*-D-glucuronidase GH67 (Glr1), and v.) Endo-xylanase GH11 (Xyn2) were PCR amplified from the generated cDNA samples using the Phusion^®^ High-Fidelity DNA polymerase (New England Biolabs) according to the manufacturer’s specification. All restriction enzymes and T4 DNA Ligase were acquired from (New England Biolabs). A complete list of plasmids, strains, and primers constructed and utilized in this study are shown in Tables S1.1 and S1.2 (Supplementary Material).

### 2.3 Plasmid Construction and Transformation

To generate the hemicellulase secretion plasmid for *S. cerevisiase*, the plasmid pBGL-AG*α*1 (a pBGLf based plasmid from Tsai, Oh et al. (2009)), utilized as a backbone for the hemicellulase secretion plasmids. The plasmid was amplified with primers pVSec_F and pVSec_R to which introduces the *BamH*I and *Not*I restriction sites downstream the *α*-factor secretion signal sequence and removes the BGL gene along with the *α*-agglutinin scaffold attached to it producing a linear fragment. The amplified hemicellulase fragments: Abf1 (1.437 kb), Axe1 (0.898 kb), Bxl1 (2.383 kb), Glr1 (2.593 kb), and Xyn2 (0.718 kb), were ligated into the *BamH*I and *Not*I sites of the linear fragment, downstream of the mating factor *α* 1 (MF*α*1) secretion sequence, generated from pBGL-AG*α*1 thus resulting into five hemicellulase secretion plasmids namely: pVSec-TrAbf1, pVSec-TrAxe1, pVSec-TrBxl1, pVSec-TrGlr1, and pVSec-TrXyn2. In the construction of the surface display strains, the C-terminal half (3’-half) of the *α*-agglutinin surface anchoring protein was amplified from the same pBGL-AG*α*1 template and ligated into the *Spe*I and *Not*I restriction sites, downstream of the hemicellulases in the pVSec plasmids, resulting into five surface display hemicellulase plasmids namely: pVSDis-TrAbf1, pVSDis-TrAxe1, pVSDis-TrBxl1, pVSDis-TrGlr1, and pVSDis-TrXyn2. These plasmids were then transformed into *S. cerevisiae* EBY100 using the standard lithium acetate transformation method as described by Bergkessel and Guthrie (2013) and selection was done in SMD+CAA media with tryptophan (TRP^−^) autotrophy.

For the creation of xylose utilizing strains, genes from the xylose isomerase pathway were utilized; as this pathway does not produce a xylitol by-product which makes this pathway quite efficient when compared to the xylose-reductase pathway and it requires minimal metabolic engineering to convert xylose into xylulose, an intermediate for the pentose-phosphate pathway (Kuyper, Winkler et al. 2004). Thus, codon-optimized xylose isomerase (XI) and xylulokinase (XK) genes from *Prevotella ruminicola* for *S. cerevisiae* were amplified from the plasmids pRH 384 and 385 (courtesy of Dr. R.Hector, USDA-ARS) (Hector, Dien et al. 2013) along with the promoter and terminator regions flanked by *Pac*I and *Asc*I sites, upstream and downstream, respectively for the XI-containing fagrment, *Asc*I and *Pm/*I sites, upstream and downstream, respectively for the XK-containing fagrment. The two amplified fragments were then cloned into the *Pac*I and *Pml*I sites of the vector-amplified fragment from the yeast vectors pXP622 (courtesy of Dr. N. Da Silva, UC Irvine) (Fang, Salmon et al. 2011), respectively to generate the plasmid pXP622-XI-XK which contains both the XI and XK genes. This plasmid was transformed into surface display EBY100 strains using the standard lithium acetate transformation method as described by Bergkessel and Guthrie (2013) and selection was done in minimal medium SMD for leucine (LEU^-^) and tryptophan (TRP^-^) autotrophy.

### Expression, Secretion, and Concentration of Hemicellulases by S. cerevisiae

*S. cerevisiae* strains harbouring the hemicellulase secretion plasmids were pre-cultured at 30°*C* for 16 hours in SMD+CAA medium, inoculated to an *OD*_600_ of ~0.5 into the 200mL of the same medium. The yeast cells were then grown at ~25°*C* and ~36h for enzyme expression and secretion. The yeast culture was centrifuged at 5000*g*, 25°*C* for 10min to separate the cells. The culture supernatant was further clarified by filtration using a 0.45*μ*m membrane filter discs (Millipore, MA, USA). Then, the clarified supernatant was concentrated ~30–50 times using Amicon^®^ Ultra-15 Centrifugal Filter unit with 30kDa molecular weight cut-off, recovered and resuspended in Reaction Buffer (50 mM citrate buffer, pH 6.0, 12 mM CaCl2, 2 mM EDTA), and stored in 4°*C*.

### 2.5 Protein Characterization

SDS-PAGE was performed via the Laemmli method, as described by Brunelle and Green (2014), using a 12% (w/v) acrylamide gel with Tris-glycine buffer (pH ~8) with 0.1% (w/v) SDS. A middle range molecular weight marker (Protech Technology Enterprise Co., Taipei, Taiwan) was utilized as a molecular weight standard.

The expressed hemicellulases were then verified via western blotting. After the SDS-PAGE, the gel was electroblotted onto a polyvinylidene fluoride (PVDF) membrane as described by Alegria-Schaffer (2014). The specific hemicellulases that were fused with a C-terminal His_6_-tag were probed with a 1°-Anti-6X His tag mouse antibody (Genetex Inc., CA, USA) then indirectly labelled by 2°-alkaline phosphatase conjugated goat Anti-Mouse antibody (Jackson Immuno Research Inc., PA, USA). The probed hemicellulases were then visualized via the alkaline phosphatase reaction using a BCIP/NBT chemiluminescent substrate (AMRESCO, OH, USA). After electroblotting, the blot was characterized using a UVP gel imaging system (UVP Bioimaging Systems, CA, USA). (Table S2.1, Supplementary Material)

The display of the hemicellulases on the surface of the surface-display strains were verified via immunofluorescence microscopy. The yeast samples (OD600nm of ~1–2) were pelleted then re-suspended in 250 μL of PBS containing 1mg/mL bovine serum albumin and 0.5 μg of 1°-Anti-6X His tag mouse antibody (Genetex Inc., CA, USA) for 4 h with occasional mixing. Then, the probed cells were pelleted, washed with PBS and resuspended in PBS plus 1 mg/ml bovine serum albumin and 0.5 μg anti-mouse IgG conjugated with Alexa 488 (Thermoscientific, MA, USA) for labelling. After incubation for 2 h, the labelled yeast cells were pelleted, washed twice with PBS, and resuspended in PBS to an OD600nm of ~1. Five µL of the labelled cell suspension were spotted onto glass slides and were further analyzed under an Olympus ix73 immunofluorescence microscope (Olympus Scientific Solutions, MA, USA). ([Figure S2.1, Supplementary Material)

### Protein Assay

The amount of enzyme present in the concentrated enzyme samples was quantified via the Bradford assay method (Bio-Rad Laboratories Inc, CA, USA) using bovine serum albumin as a standard. The amount of enzyme was expressed in terms of mg BSA equivalents.

### 2.7 Enzyme Activity Assay

The activity of the hemicellulases secreted by *S. cerevisiae* was assayed by determining the reducing sugar release rate upon the hydrolysis of the hemicellulosic substrates, beechwod xylan (Sigma, USA) and Wheat Arabinoxylan (low viscosity; Megazyme, USA). The assay was carried out as follows in accordance to the method described by Vazana, Moraïs et al. (2012): to 100*µ*L of enzyme sample (pure or mixture) in reaction buffer, 50*µ*L of 3% xylan or 100 *µ*L 1.5% Arabinoxylan solutions (in 50mM citrate buffer, pH 6) was added to commence the reaction and was continued for 45 minutes at 30°*C*. The reaction was stopped by transferring the tubes to an ice-water bath. The amount of reducing sugars released in the reaction was determined via the DNS method described by Wood and Bha (1988) using xylose as a standard. One unit of enzyme activity (U) was defined as the amount of xylose released (*µ*mol) per 1min under assay conditions.

### 2.8 Mixture Experimental Design Setup

With the aid of the Minitab^®^ Statistical Software (Minitab 2014), a five(5)-component, augmented simplex lattice design, with a lattice degree of 3 was generated and is summarized and visualized in a triangular coordinate plot in Table S3.1, and Figure S3.1 (Supplementary Material). This was adapted to investigate the synergistic relationships of the five hemicellulases secreted by *S. cerevisiae* in their capability to degrade the xylan substrate. The response data was analysed using the said statistical software. Xylan hydrolysis for each mixture formulation were performed in triplicates.

### 2.9 Mixed Culture Fermentation Experiments

After pre-cultivation, the different yeast cells expressing the xylose utilization genes along with a respective surface-display hemicellulase were harvested, washed, and resuspended with 50mM Na-citrate buffer (~pH 6) containing 12mM *CaCl_2_* and 2mM EDTA. A mixture of these yeast cells which corresponds to their respective optimum hemicellulase mixture formulation were then inoculated into synthetic medium containing 1–2% Beechwood xylan or 1–2% Wheat Arabinoxylan at a total initial *OD*_600_ was fixed at ~3.0–5.0. Ethanol fermentation was performed in shake-flasks at 30°C and 150rpm. All fermentations were performed in duplicates.

Yeast growth monitoring was estimated by measuring the *OD*_600_ using a Synergy H1 microplate reader (Biotek, VT, USA) corrected with the *OD*_600_ values of the hemicellulose media as blanks. Ethanol and acetic acid were detected via gas chromatography on a GC2010 (Shimadzu Scientific Instruments, Taiwan) equipped with a flame ionization detector (FID) along with a Stabilwax-DA (Restek, PA, USA) capillary column.

## 3 Results

### 3.1 Mixture Design Experiments

For the efficient release of xylose monomers from the xylan backbone, at least five different types of hemicellulases are needed (Moreira and Filho 2016). Thus, a mixture experimental design approach was applied to describe the overall mixture behaviour of these hemicellulases when exposed to two different xylan substrates (beechwood xylan and wheat arabinoxylan) with different accessibilities (based on the difference in viscosities of these substrates). Contour plots for the mixture design are shown in Figure 2.

**Figure 2.**
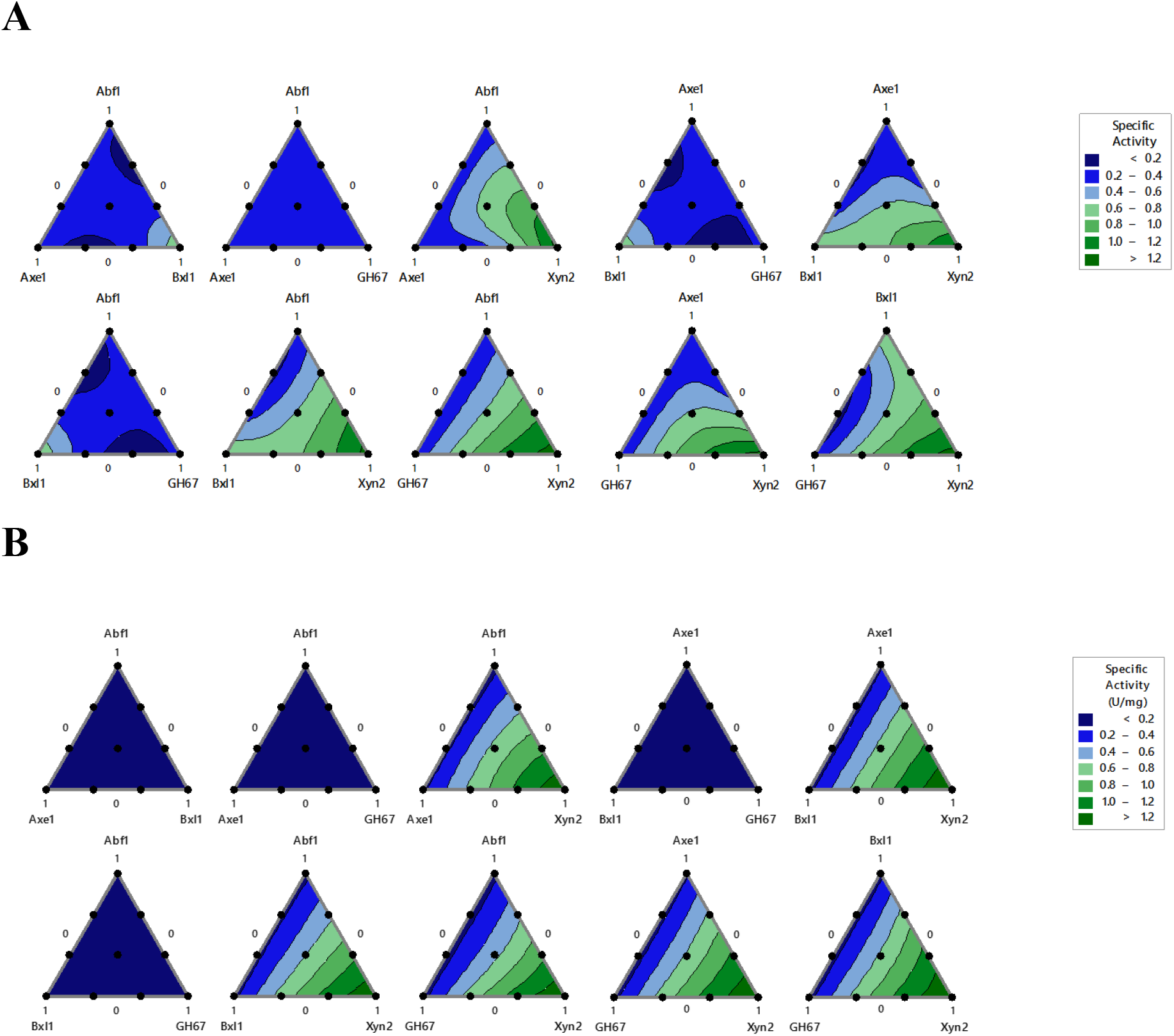
Minitab^®^ generated Mixture Contour Plots for the Specific Activity: Beechwood Xylan Hydrolysis **(A)**, and Wheat Arabinoxylan Hydrolysis **(B)**.

#### 3.1.1 Hemicellulase Mixture Optimization for Xylan Hydrolysis

The specific activities of the different hemicellulases mixtures on the hydrolysis of beechwod xylan substrate are described as Contour plots and presented in Figure 2A. The model summary statistics are shown in Tables S3.4 and S3.5 (Supplementary Material). The specific activities of the different hemicellulase mixtures on beechwood xylan hydrolysis are adequately described by a cubic model Table S3.4; which implies that the three-component synergistic interactions are significant. These three-component synergistic interactions (blending/synergism terms or the polynomial coefficients of the mixture model) are schematically illustrated in Figure 3. It is evident that high specific activities and synergistic interactions are observed when Xyn2 is a major component in the mixture; as it is one of the main-chain cleaving enzymes in xylan degradation. The main-chain cleaving hemicellulases, Xyn2 and Bxl1, are positively complemented with the following accessory hemicellulases: Axe1 (with a blending synergism value of ~6.8), Abf1 (~3.5). Furthermore, the highest synergistic interaction obtained from this experiment exists between Abf1, Axe1, and Xyn2 (with a blending synergism of ~8.5).

**Figure 3.**
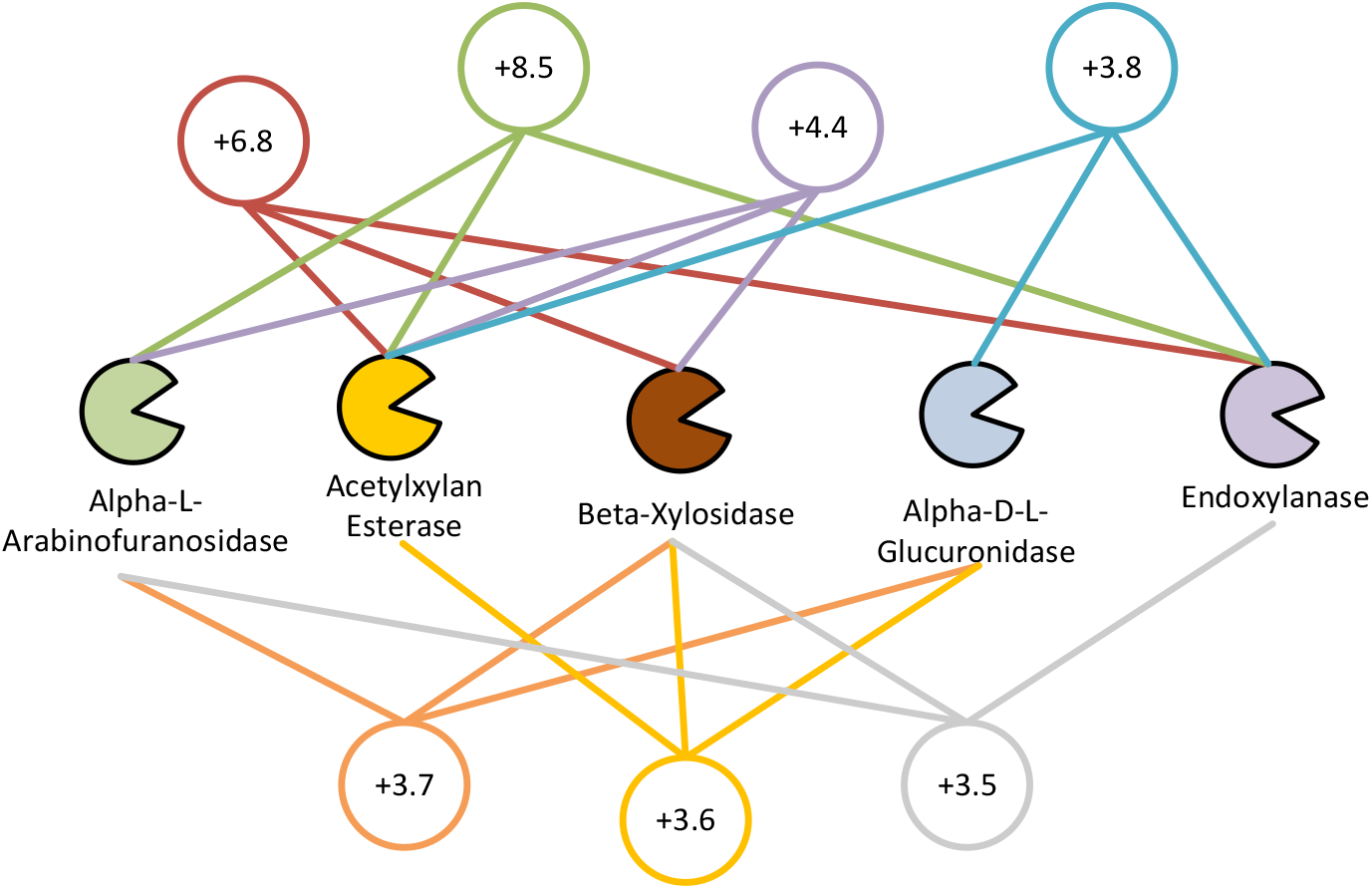
Ternary component interaction between hemicellulases

For the efficient release of xylose monomers from the xylan backbone, the condition that all the five hemicellulases must be present in the hydrolysis mixture was adapted during the mixture response optimization process. The optimum mixture formulation for the given constraint is shown in Table 1 having a predicted specific activity of 1.104 U/mg with a ~8% difference when compared to the experimentally obtained activity; which implies the adequacy of the model.

**Table 1.** Predicted and Experimental Specific Activities

| Substrate | Mixture Composition<br>(Mass Fractions) |  |  |  |  | Specific Activity (U/mg) |  |  |
| --- | --- | --- | --- | --- | --- | --- | --- | --- |
|  | Abf1 | Axe1 | Bxl1 | Glr1 | Xyn2 | Predicted | Experimental | Percent<br>Difference<br>(%) |
| Beechwood<br>Xylan | 0.14 | 0.16 | 0.11 | 0.12 | 0.47 | 1.143 | 1.197 | 8.42 |
| Wheat<br>Arabinoxylan | 0.10 | 0.10 | 0.10 | 0.10 | 0.60 | 0.834 | 1.032 | 23.74 |

#### 3.1.2 Hemicellulase Mixture Optimization for Arabinoxylan Hydrolysis

The specific activities of hemicellulase mixtures on the hydrolysis of wheat arabinoxylan substrate are described as Contour plots in Figure 2B and the corresponding model summary are shown in Tables S3.6 and S3.7 (Supplementary Material). In contrast to the hydrolysis of beechwood xylan, the variations in the specific activities (response) with different hemicellulase mixture formulations are not adequately described by higher ordered models which in this case suggests that the linear terms are sufficient to predict the response. In other words, the synergistic interactions between the hemicellulases during arabinoxylan hydrolysis are not statistically significant. As illustrated in the Countour plot, it would be sufficient to say that hydrolysis of the substrate by using the main-chain cleaving enzyme, Xyn2, is necessary prior to the addition of the other hemicellulases.

Thus the mixture formulation that is shown in Table 1 was chosen under the condition that all the five hemicellulases be present for efficient xylose release from the xylan backbone. The difference of ~24% between the predicted and experimental specific activities thus supports the inadequacy of the model to explain the variations due to the hemicellulase synergies. This mixture formulation was adapted to further investigate on the ethanol fermentation from arabinoxylan.

### 3.2 Hemicellulose Hydrolysis Experiments by Whole-Cell Biocatalysts Mixtures

It was to the understanding that the overall specific activities of the hemicellulase mixtures for xylan hydrolysis can be enhanced by displaying the hemicellulases on the yeast surface to serve as whole-cell biocatalysts. Thus, each hemicellulase was displayed on the surface of yeast and its mixture formulations (which corresponds to the free hemicellulase mixture formulation state in Table 1 were evaluated for its overall hydrolysis activities on xylan substrates. Specifically the hydrolysis activities of the following systems were evaluated on different xylan substrates: i.) the free hemicellulase mixture formulation obtain from the mixture of single strain secreted hemicellulase isolates Figure 4A), ii.) mixed culture secreted hemicellulase formulation Figure 4B), and iii.) mixed culture surface display formulation Figure 4C).

**Figure 4.**
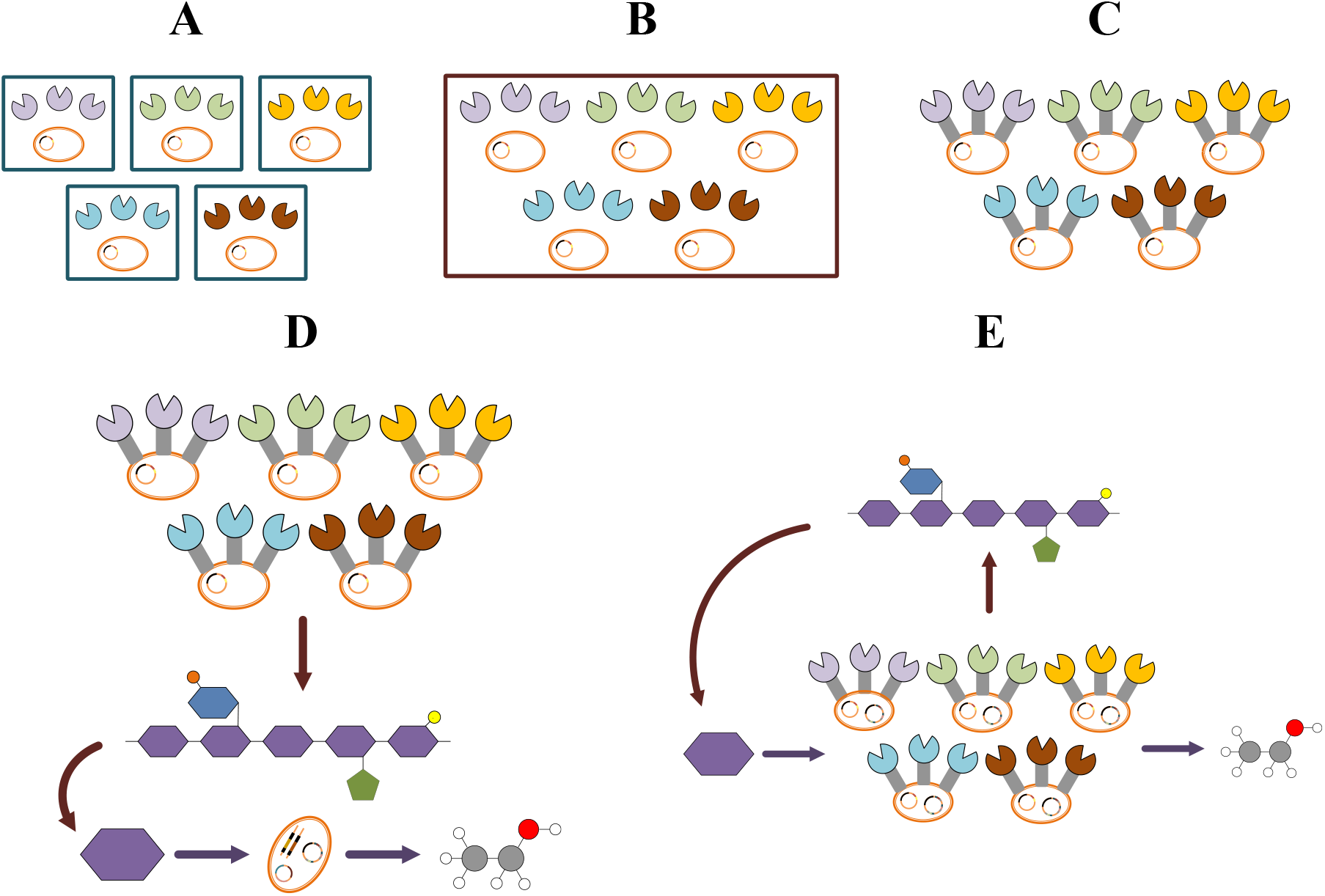
Single and Mixed Culture Systems employed in this study. Single strain enzyme secretion isolate system **(A)**, Mixed culture secretion enzyme isolate system **(B)**, Mixed culture surface display system **(C)**, Mixed culture fermentation system containing hemicellulase-displaying yeast strains along with a xylose-utilizing yeast YRH1114 **(D)**, Mixed culture fermentation system containing hemicellulase-displaying, xylose-utilizing yeast strains **(E)**.

The specific activities of these different mixed culture system for different xylan substrates are summarized in Table 2 It is evident that the specific activities of the surface-display hemicellulase formulation have enhanced the free hemicellulase mixture activities on beechwood xylan and wheat arabinoxylan hydrolysis by ~70 and ~25% respectively.

**Table 2.** Hydrolysis of Xylan Substrates by Whole-Cell Biocatalyst Mixed Cultures

| Substrate | Mixture Composition<br>(Mass Fractions) |  |  |  |  | Specific Activity (U/g cell) |  |  |  |
| --- | --- | --- | --- | --- | --- | --- | --- | --- | --- |
|  | Abf1 | Axe1 | Bxl1 | GH67 | Xyn2 | Free<br>Enzyme<br>Mixture<br>[A] | Free<br>Enzyme<br>Mixed<br>Culture<br>[B] | Cell-Surface<br>Display<br>Mixture<br>[C] | Fold<br>Change<br>[C] to [A] |
| Beechwood<br>Xylan | 0.14 | 0.16 | 0.11 | 0.12 | 0.47 | 0.342 | 0.426 | 0.570 | 1.67 |
| Wheat<br>Arabinoxylan | 0.10 | 0.10 | 0.10 | 0.10 | 0.60 | 0.305 | 0.314 | 0.382 | 1.25 |

### 3.3 A Simple Growth Assay for the Simultaneous Xylose Release and Utilization during Xylan Hydrolysis

To assess the quality of the xylan hydrolysis products (release of xylose in the medium) by the mixed culture surface display system, a simple xylose growth assay was adapted. Specifically, to a hemicellulase surface display mixed culture formulation, a xylose-utilizing “control” strain YRH1114 (adapted for improved xylose utilization, (Hector, Dien et al. 2013)) was added to assess that the xylose released by the hemicellulase surface display mixture can be utilized by the strain YRH1114 to produce ethanol.

Thus, from the results shown in Figure 5A, there is an observable cell growth associated to the increase in the 0D600 measurements (which used as an estimate for cell concentration; corrected with the absorbance readings of the xylan media as blank). The cell growth observed in Figure 5A was further supported by the released reducing sugar (RRS) profile shown in Figure 5B. In addition, the RRS profile for the hemicellulase surface display mixed culture system (Setup C, refer to Figure 4) shows higher rate of released reducing sugars as the fermentation progresses in contrast to the xylose growth assay assessment system (Setup D, refer to Figure 4). This difference in the rates of released reducing sugar supports the claim that the xylose released in the medium by surface display hemicellulase mixed culture was utilized by the control strain YRH114. Furthermore, around ~150mg/L ethanol was produced at 120hrs of the fermentation experiment as shown in Figure 5C as a product of xylose utilization by the xylose-utilizing control strain.

**Figure 5.**
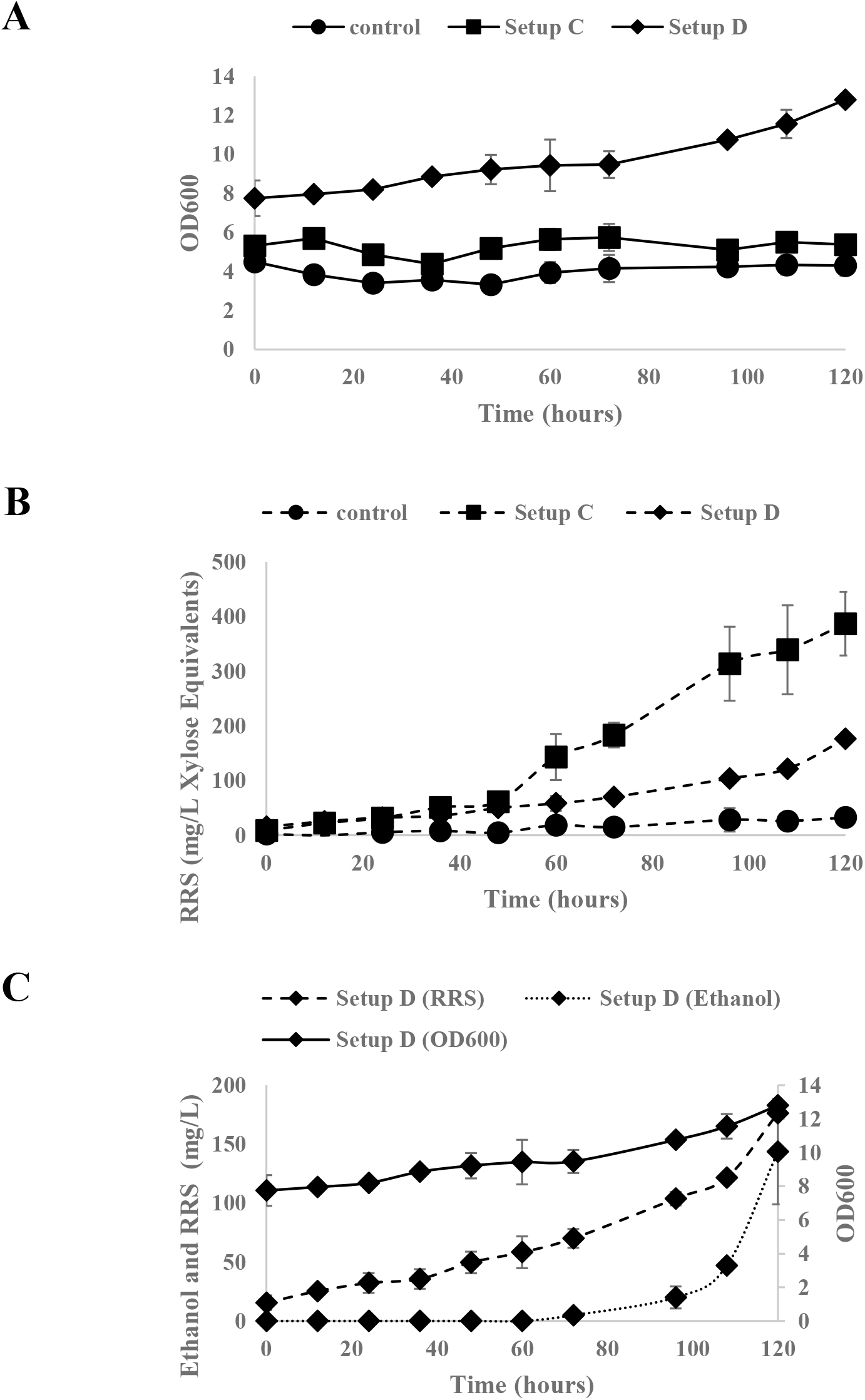
Xylose Growth Assay on the Degradation and Utilization of 2% beechwood xylan in minimal media by the mixed cultivation system C and D. Fermentation Profiles: OD600 **(A)**, Released Reducing Sugars (RRS) expressed in xylose equivalents **(B)**, OD600, RRS, and Ethanol Profiles of Mixed cultivation system D **(C)**. Presented data are averages from duplicate cultivation experiments and Error bars represent the corresponding standard deviations

### 3.4 Ethanol Fermentation of Xylan Substrates by Hemicellulase-Displaying, Xylose-Utilizing Yeast Strains

As the hydrolysis products of the hemicellulase surface-display mixed cultures can be utilized by a xylose-utilizing yeast strain, the xylose isomerase (XI) and xylulokinase (XK) genes from the xylose-utilizing strain were then cloned into each of the hemicellulase surface-display strains to construct the respective hemicellulase-displaying, xylose-utilizing yeast strains. These resulting strains were able to grow on xylose as a sole carbon source in a minimal medium at specific growth rates ranging from ~0.02 to ~0.03 hr^−1^ (Table S4.1 and Fig. S4.1, Supplementary Material). It was noted that during the fermentation media preparation, the wheat arabinoxylan medium (WAM) was slightly viscous with regards to the beechwood xylan medium (BXM).

The ethanol fermentation profiles of xylan substrates by engineered yeast mixed cultures (whose formulations correspond to the free enzyme formulations for xylan substrates shown in Table 1) are presented in Figure 6. It can be observed that the overall mixed culture growth on BXM is lower than that of the growth on WAM. This difference in specific growth rates of the mixed cultures between the BXM and WAM are supported by the release rates of the reducing sugars (RRS). Additionally, it can be inferred that the simultaneous rate of release and consumption of xylose in WA medium is higher than that in the BX medium; which could be related to the hemicellulase accessibility on the different substrates.

**Figure 6.**
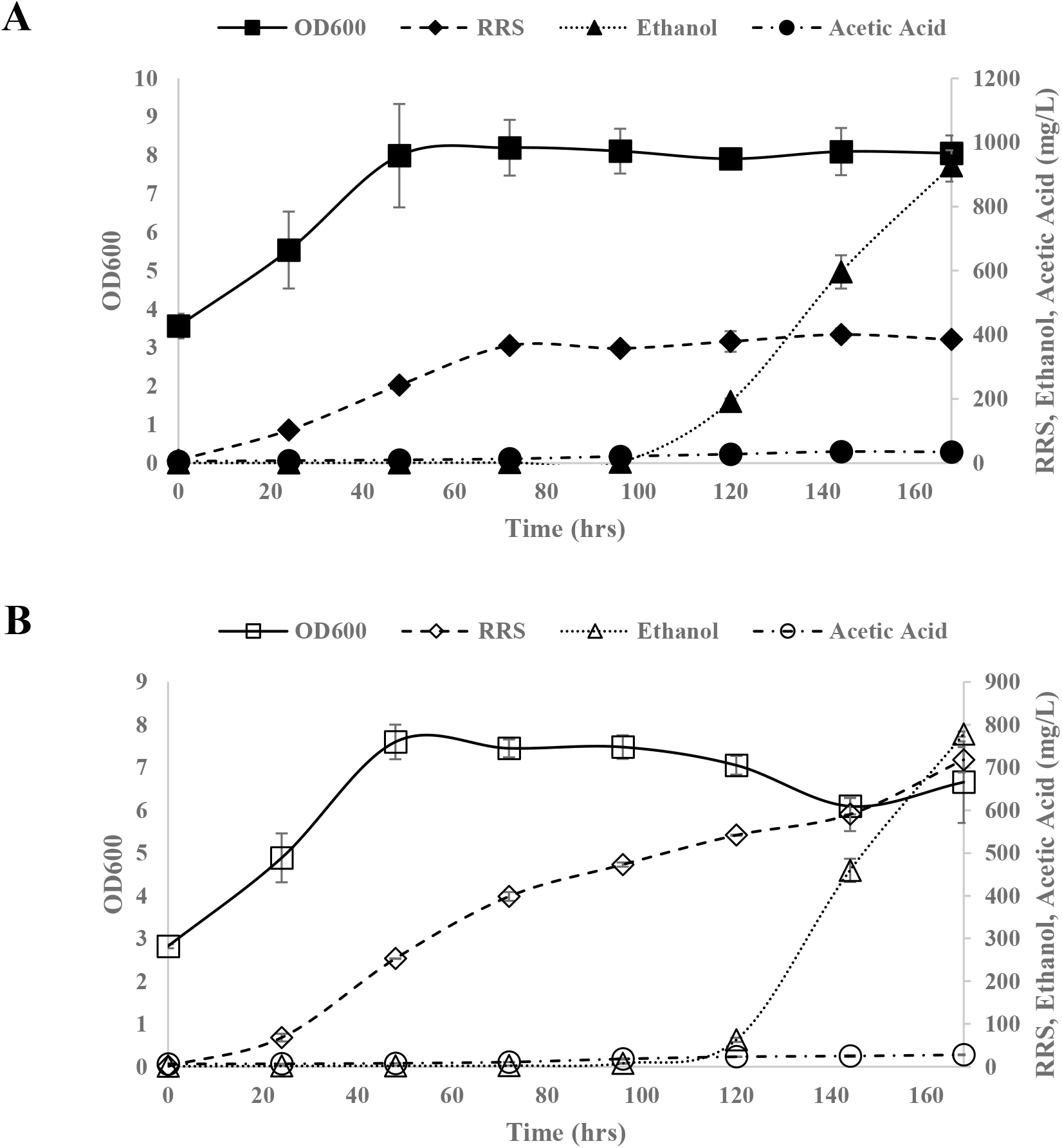
Fermentation profiles of the mixed cultivation system E for the following hemicellulose substrates in minimal media: 1% Beechwood Xylan **(A)**, 1% Wheat Arabinoxylan **(B)**. Presented data are averages from duplicate fermentation experiments and Error bars represent the corresponding standard deviations

The fermentation parameters for both BX and WA substrates are summarized in Table 3. The determined percentage of ethanol obtained from the fermentation experiment with respect to the theoretical ethanol yield from xylose for both BX and WA substrates are ~22 and ~25%, respectively.

**Table 3.** Fermentation Parameters of Xylan Substrates by Engineered Yeast Mixed Cultures

| Xylan Substrates | Beechwood Xylan |  | Wheat Arabinoxylan |  |
| --- | --- | --- | --- | --- |
|  | Average | SD | Average | SD |
| **Specific Growth Rate $\mu$ (hr<sup>-1</sup>)** | 0.0167 | 0.0016 | 0.0206 | 0.0007 |
| **Ethanol Titer (mg/L)** | 960.97 | 0.51 | 778.13 | 6.14 |
| **Acetic Acid Titer (mg/L)** | 34.72 | 1.78 | 27.92 | 0.99 |
| **% Theoretical EtOH Yield <sup>[1]</sup>** | 21.55 | 0.01 | 24.54 | 0.19 |

[1] –% Theoretical ethanol yield is determined based on the amount of ethanol obtained experimentally with respect to the amount of ethanol produced if all the xylose in the substrate were converted to ethanol [based on the $Y_{PS}$ of 0.51g ethanol per g xylose (Kuyper, Winkler et al. 2004)]
\* - all fermentations were performed in duplicates
\*\* -SD values represent standard deviation

## 4 Discussion

### 4.1 Mixture Design Experiments

In studying the interactions of mixture components, the mixture experimental design methodology is an indispensable tool for modelling and optimization of the response variables with respect to varying mixture formulations. Not only that the optimum formulation to produce the desired (maximum or minimum) response be obtained, it can also provide the blending properties (synergism or interaction) between the mixture components (Cornell 2011). Therefore, this methodology was adapted in this experiment to obtain a statistical understanding on the mechanisms involved in the hydrolytic activities of the five different hemicellulase mixtures expressed and secreted in *S. cerevisiae* strains on xylan substrates. Additionally, this design approach has been utilized for determining optimum formulations of hemicellulase mixtures and also, along with cellulases (either commercially or recombinantly prepared) in the hydrolysis of not only pure polysaccharide substrates but with actual lignocellulosic substrates (e.g. rice straw, sugarcane bagasse, etc.) (Sørensen, Pedersen et al. 2005, Berlin, Maximenko et al. 2007, Suwannarangsee, Bunterngsook et al. 2012, Goldbeck, Damásio et al. 2014, Laothanachareon, Bunterngsook et al. 2015).

In the hydrolysis of beechwood xylan substrate, homo-synergy between main-chain cleaving enzymes (Bxl1 and Xyn2) was complemented with the activities of the accessory or debranching enzymes (Abf1, Axe1, although no further synergistic effect in the addition of Glr1). These synergistic interactions are commonly observed in hydrolysis studies of xylan from birchwood, beechwood, and oat-spelt xylan (which are considered as relatively soluble xylans) using hemicellulases from various fungal microorganisms (e.g. *Trichoderma reesei, Aspergillus niger, Clostridium thermocellum, etc*.) (Gasparic, Martin et al. 1995, Raweesri, Riangrungrojana et al. 2008, Carapito, Carapito et al. 2009, Shi, Li et al. 2010, McClendon, Shin et al. 2011).

On the other hand, the mixture design methodology for the hydrolysis of wheat arbinoxylan did not yield significant synergistic interactions between the hemicellulases. Rather, the observed enhanced degree of synergy values (not to be confused with the blending synergism obtained from the fitted mixture model) suggest that some extent of synergism is observed when there is a significant amount of main-chain cleaving hemicellulases such as Xyn2 and Bxl1 are present in the hemicellulase mixture (Table S3.3, Supplementary Material). This case was also observed in the studies of Sorensen et al., where the release of xylose is due to the initial activity of the endoxylanase which is then further supplemented with the *β*-xylosidase activity (Sørensen, Meyer et al. 2003, Sørensen, Pedersen et al. 2005, Sørensen, Pedersen et al. 2006, Sørensen, Pedersen et al. 2007).

### 4.2 Whole-Cell Biocatalysts for the Hydrolysis of Xylan Substrates

The application of yeast cell surface display on to the hydrolysis of xylan substrates provides a platform where the hemicellulases are immobilized on the cells and can provide enhancements of their enzymatic activities, localized enzyme concentration on the yeast surface (which relates to higher enzyme loading), operational stability, and recyclability (Kondo and Ueda 2004, Liu, Ho et al. 2016, Ueda 2016, Grunwald 2017). Furthermore, in achieving control of the hemicellulase mixture formulations, a mixed culture strategy was adapted and from the study of Baek, Kim et al. (2012); which is also applied into yeast consortium studies for consolidated bioprocessing (Tsai, Goyal et al. 2010, Goyal, Tsai et al. 2011, Fan, Zhang et al. 2012, Kim, Baek et al. 2013).

Thus the experimental data from the surface-display hemicellulase mixture formulations support the claim that the enhancement of the specific activities on the hydrolysis of xylan substrates is associated to the immobilization of the hemicellulases on the yeast surface.

### 4.3 Growth Assay for the Simultaneous Xylose Release and Utilization during Xylan Hydrolysis

The adapted growth assay for assessing the utilization of xylan hydrolysis products by the mixed culture surface display system was inspired from symbiotic co-culture studies on efficient utilization of lignocellulosic substrates for consolidated bioprocessing (Panagiotou, Topakas et al. 2011, Zuroff and Curtis 2012, Brethauer and Studer 2014). Specifically, the main idea for this symbiotic co-culture strategy is that one microorganism (with good substrate utilization ability) is responsible for the efficient substrate breakdown and another (with good product yield) for the utilization of the deconstructed substrate. However in this experiment, instead of using two different symbiotic microorganisms to achieve xylan hydrolysis and xylose utilization, two sets of yeast strains were employed where one strain (referred as the “control” strain) is capable of xylose utilization and the other set would be a group of strain/s that express hemicellulases for efficient xylan degradation. Thus, the utilization of a xylose-utilizing strain to test if the xylan degradation products contain xylose provides a simple alternative in characterizing the efficiency of xylan hydrolysis for fermentation experiments when compared to the tedious and costly chromatographic separation techniques.

### 4.4 Ethanol Fermentation of Xylan Substrates

The observed differences in the overall growth, and simultaneous xylose release and consumption of mixed cultures in BXM and WAM can be attributed to the accessibilities of the whole-cell biocatalysts to their respective substrates. Specifically, it can be inferred that for there should be a better hemicellulase accessibility to the WA substrate, which is directly related to the specific activity and growth of the mixed cultures, when compared to those in the BX substrate. One factor that could affect the accessibility of the hemicellulases on the substrates would be the viscosity (Sørensen, Pedersen et al. 2006). As it was observed that the WAM has a higher viscosity when compared to BXM; owing to the inherent property of arabinoxylan to produce a viscous solution when dissolved in the media (Andrewartha, Phillips et al. 1979, Courtin and Delcour 2001, Courtin and Delcour 2002). Furthermore, due to the viscous nature of WAM, the yeast cells displaying hemicellulases have a better dispersion in the medium and thus providing a good access for the yeast cells to release and utilize xylose from the substrate.

The different studies on xylan utilization by *S. cerevisiae* are summarized in Table 4 and this shows that the hemicellulase surface display mixed culture strategy employed in this study for the degradation and utilization of xylan substrates exhibit a comparable performance to those strains that are adapted for xylose utilization (based on the specific growth rates and % theoretical yield values). However, current engineered strains designed for ethanol fermentation of xylan substrates exhibit low yields when compared to their starch and cellulose counterparts and thus provides a potential for improvement in this field (Schuster and Chinn 2013, Hasunuma, Yamada et al. 2014, Favaro, Viktor et al. 2015, Haan, Rensburg et al. 2015, Sigoillot and Faulds 2016, Ueda 2016, Ábrego, Chen et al. 2017, Chen 2017).

**Table 4.**
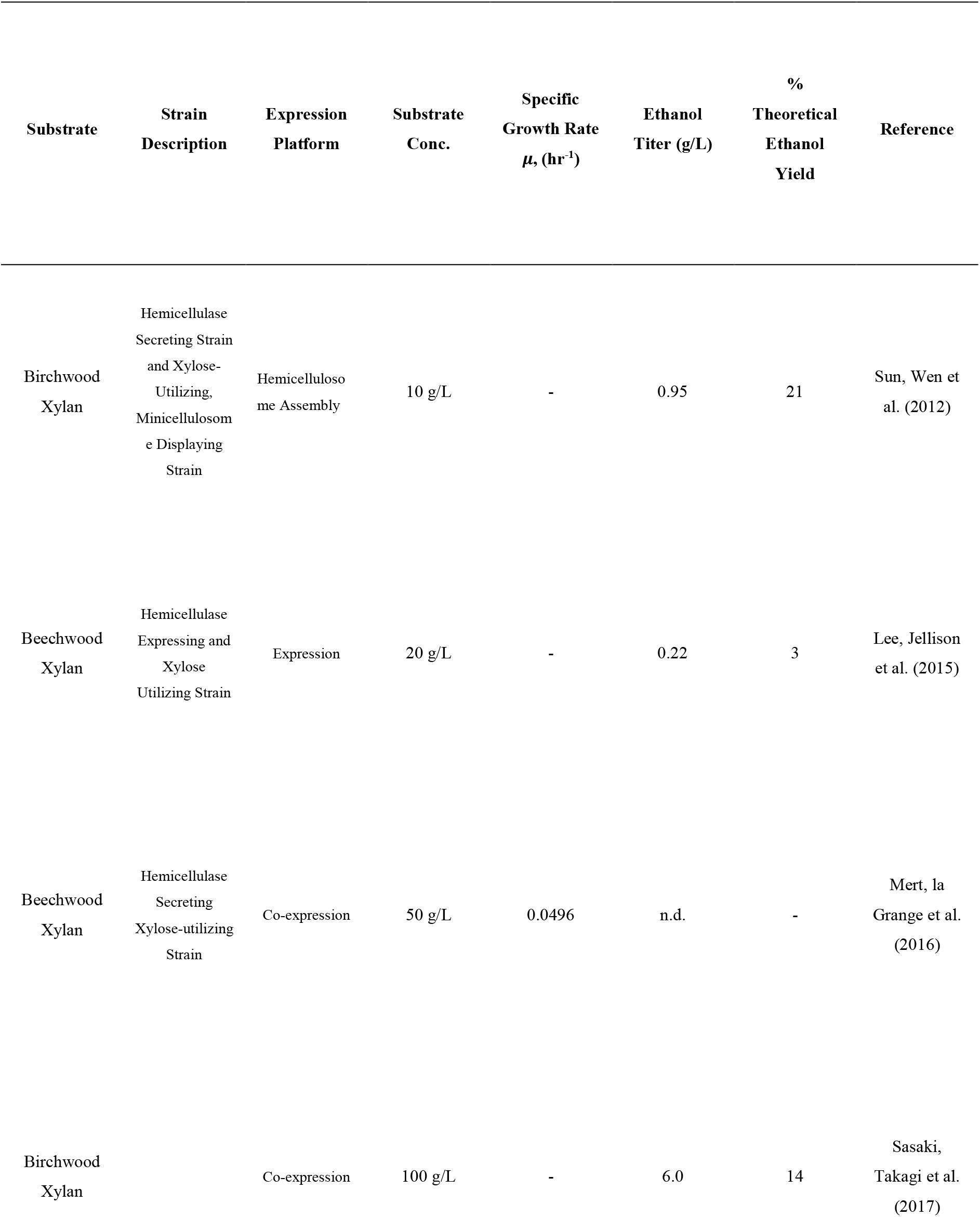
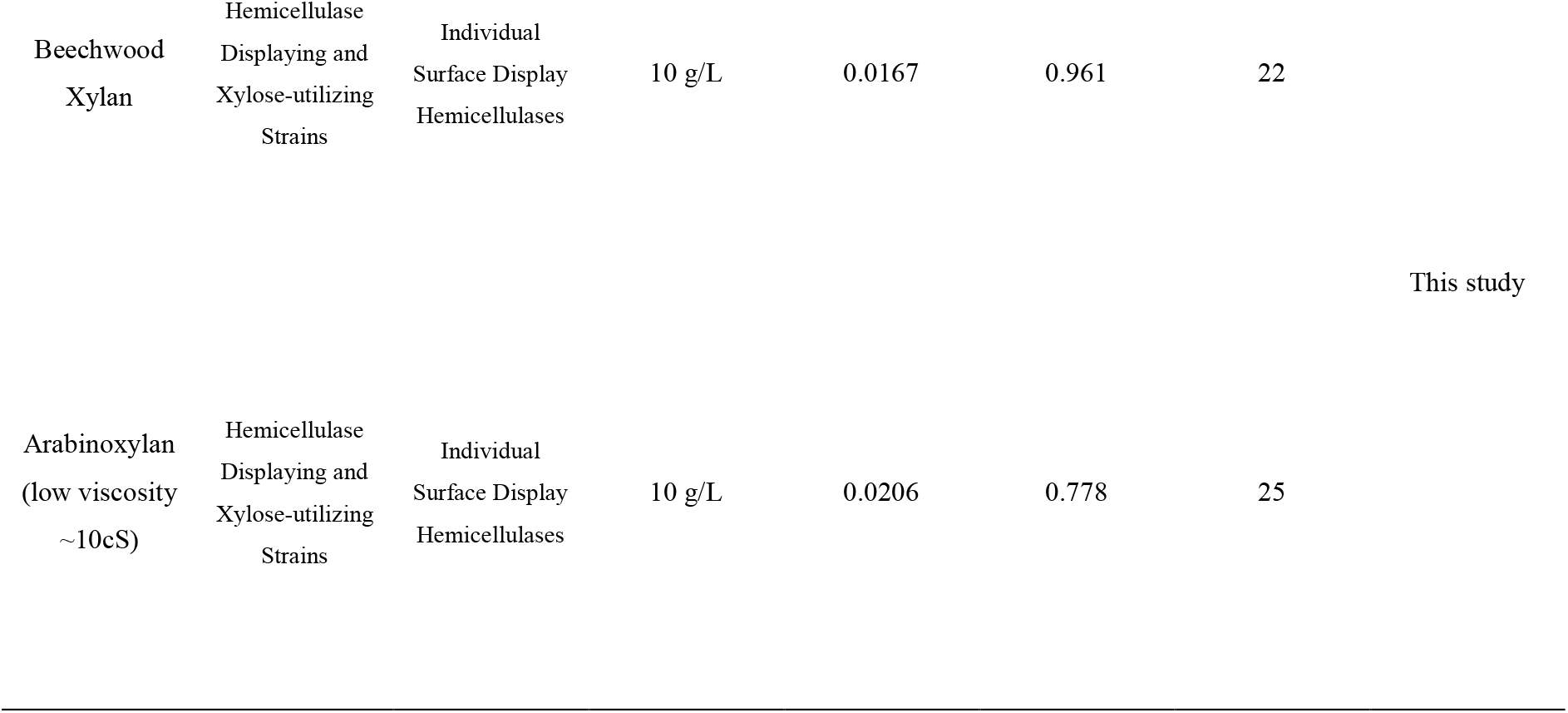
Comparisons on the different studies on Xylan Utilization by Engineered Yeasts

## 5 Conclusions

This study provides an insight into the different statistically significant hemicellulase interactions in the hydrolysis of xylan substrates by employing the mixture design experiments. Then the succeeding experiments demonstrates that the xylan hydrolysis activities are enhanced by displaying the hemicellulases on the yeast surface to produce whole-cell biocatalysts. Furthermore, the adapted xylose growth assay in characterizing the release of xylose in the xylan hydrolysis products provides a good alternative in assessing future xylan hydrolysis experiments. Finally, this is a first demonstration on application of the mixed culture strategy in controlling the mixture formulations of different hemicellulase-displaying xylose-utilizing yeast strains for the efficient degradation and utilization of xylan substrates.

